# Persister Cells Form Based on Low Ribosome Content

**DOI:** 10.64898/2025.12.18.695105

**Authors:** Hyeon-Ji Hwang, Rodolfo García-Contreras, Michael J Benedik, Thomas K. Wood

## Abstract

Persister cells survive any severe stress including antibiotics, starvation, heat, oxidative conditions, and phage attack, by entering a dormant physiological state. They arise without genetic change and can resume growth once the stress is removed and nutrients are available. Critically, upon resuscitation, persister cells can reconstitute infections. Although it is known that persister cells *resuscitate* in proportion to their ribosome content, it has remained unclear whether ribosome levels also influence the *formation* of persister cells. Here, we used fluorescence-activated cell sorting (FACS) to fractionate exponentially growing cells into four populations spanning low to high ribosome levels and demonstrated that cells with low ribosome content form persister cells approximately 80-fold more frequently than cells with population-average ribosome levels. These findings show that persister cell formation is inversely proportional to ribosome abundance. Cells with low ribosome levels are less metabolically-active and therefore less capable of initiating a stress response like most cells; instead, they become dormant.

## INTRODUCTION

It is estimated that treating bacterial infections worldwide will cost $100 trillion per year in 2050 (Udaondo and Matilla, 2020). Currently, about 14 million (Ikuta et al., 2022) to 17 million (Martens and Demain, 2017) die each year from bacterial infections, and these infections are the second leading cause of death worldwide. Although millions of lives are saved each year from antibiotics (Martens and Demain, 2017), it was discovered at the beginning of their widespread use, that a small population of bacterial cells (less than one percent) survive antibiotic treatment (Hobby et al., 1942); these surviving cells were termed ‘persisters’ (Bigger, 1944). Persister cells do not form spontaneously (‘stochastically’), but, instead, form as a result of elegant regulation in response to stress (Song and Wood, 2020a). They also do not resuscitate spontaneously, but instead, revive based on the presence of nutrients in the absence of stress (Yamasaki et al., 2020; Fang and Allison, 2023). Critically, most forms of lethal stress induce persistence, including antibiotics (Kwan et al., 2013), oxidative conditions (Hong et al., 2012), acid conditions (Hong et al., 2012), ultraviolet light (Ichikawa et al., 2022), nutrient limitations (Kim et al., 2018a) and bacteriophages (Fernández-García et al., 2024).

By chemically-inducing dormancy by converting the whole population of exponentially-growing *Escherichia coli* cells into persister cells by halting either transcription or translation (Kwan et al., 2013) and by studying single cells (Kim et al., 2018b; Song et al., 2019; Song and Wood, 2020a, b; Yamasaki et al., 2020; Song et al., 2021), it was discovered that persister cells form by inactivating their ribosomes through the alarmone/stress signal guanosine pentaphosphate/tetraphosphate (henceforth, (p)ppGpp). (p)ppGpp halts translation through ribosome-associated inhibitor (RaiA) (via ribosome inactivation), ribosome modulation factor (RMF), and hibernation promoting factor (Hpf) (via ribosome dimerization); this persister formation/resuscitation mechanism is termed the ‘ppGpp Ribosome Dimerization Persister (PRDP)’ model (Song and Wood, 2020a; Wood and Song, 2020). Additional features of this mechanism include that persister cells resuscitate in a heterogeneous manner based on the disparate number of active ribosomes (Kim et al., 2018b) via the high frequency of lysogenization X (HflX) (via ribosome splitting) (Yamasaki et al., 2020), with most cells waking in minutes (Kim et al., 2018b), and persister cells resuscitate based on nutrient recognition through chemotaxis proteins and the phosphotransferase system (Yamasaki et al., 2020). The PRDP model for persister cells appears general since all three domains dimerize ribosomes (Beckert et al., 2018; Yaeshima et al., 2022; McLaren et al., 2023; van Schijndel, 2023). Non-molecular competing models include the ‘Persistence as Stuff Happens’ model that posits that persister cells form due to various replication errors (Johnson and Levin, 2013), and the ‘Randomly Connected Cycles Network’ model that hypothesizes that persisters formation is not regulated and depends on a random network (Kaplan et al., 2021).

Given that persister cells resuscitate according to the number of active ribosomes (Kim et al., 2018b), we reasoned that the inverse should also be true, namely that cells with fewer ribosomes should enter the persister state more rapidly. Here, we demonstrate that the cellular ribosome level indeed dictates the rate of persister cell formation.

## MATERIALS AND METHODS

### Bacterial strains and culture conditions

*E. coli* MG1655-ASV, which expresses the green fluorescent protein (GFP) in proportion to ribosome content (*rrnB* P1::*gfp*) (Shah et al., 2006), and *E. coli* MG1655 (used as the GFP-negative control for defining FACS gating, **Fig. S1** and **S2**) were cultured in Luria-Bertani (LB) medium (tryptone 10 g/L, yeast extract 5 g/L, and NaCl 10 g/L) at 37°C with vigorous shaking at 250 rpm. For solid medium, agar was added to a final concentration of 1.5% (wt/vol). Bacterial growth was measured by optical density at 600 nm (OD_600_). Ampicillin (100 μg/mL) was used to eliminate non-persister cells (Kim et al., 2018b).

### FACS analysis and cell sorting

The initial analysis was performed using a Cytek Aurora analyzer (Cytek Biosciences, Fremont, CA). Subsequent analyses and all sorting procedures were conducted on a Bigfoot cell sorter (Thermo Fisher Scientific, Waltham, MA) at the Huck Flow Cytometry Facility. Overnight cultures grown in LB were diluted 1:500 into 20 mL of fresh LB and incubated at 37°C with shaking until reaching an OD_600_ of 1. A 1 mL aliquot was harvested by centrifugation at 13,300 × g for 6 min and resuspended in 1 mL of phosphate-buffered saline (PBS; 8 g/L NaCl, 0.2 g/L KCl, 1.44 g/L Na_2_HPO_4_, 0.24 g/L KH_2_PO_4_, pH 7.4). The suspension was then diluted 100-fold to obtain the recommended concentration for sorting (5 to 25 × 10^6^ cells/mL). Cells were first gated on forward scatter (FSC), assisted by 405 nm small-particle detection and 488 nm-based side scatter (SSC), to exclude debris and define the main population. GFP fluorescence was excited using a 488 nm blue laser and detected with a 507/19 nm bandpass filter. Subpopulations of bacterial cells were sorted in purity mode at 37°C based on GFP intensity, yielding approximately 10^5^ cells per group for the four groups (A–D) (**Fig. 1A**).

**Fig. 1.**
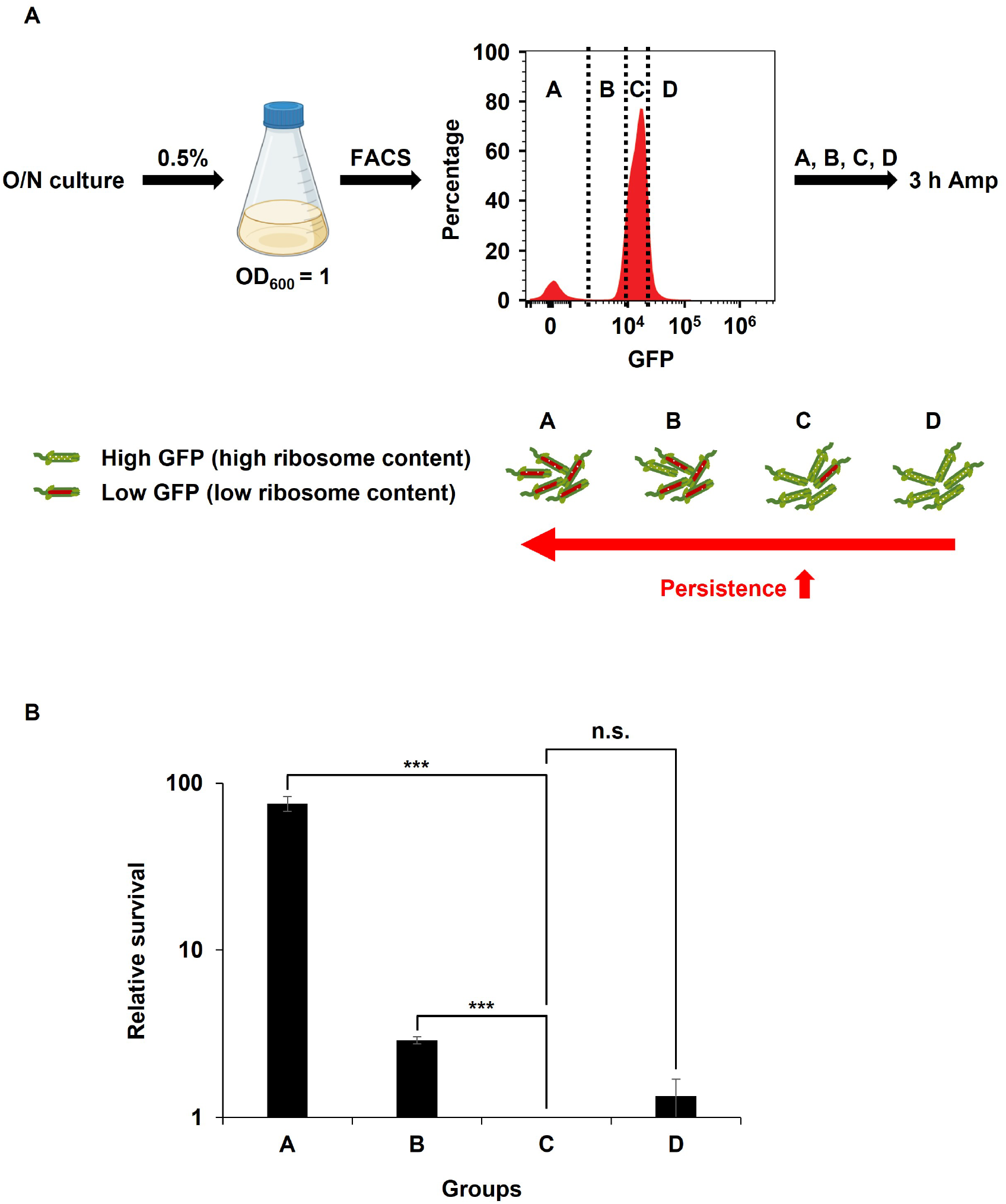
FACS-based separation of cells by ribosome levels and corresponding differences in persister frequency. (**A**) Schematic illustrating the sorting strategy used to separate *E. coli* MG1655-ASV cells into four groups (A–D) based on GFP fluorescence intensity, which reflects ribosome abundance. The schematic also conceptually illustrates the hypothesized inverse relationship between ribosome levels and persister frequency. (**B**) Relative persister frequencies of the four sorted groups. Relative survival was calculated from CFU counts normalized to Group C (set to 1). ***, *P* < 0.005; n.s., not significant.

### Persister cell viability assay

Immediately after sorting, each cell group was collected by centrifugation at 13,000 × g for 6 min and resuspended in 1 mL of fresh LB containing ampicillin (100 µg/mL, 10× MIC, (Kwan et al., 2013)) in a 15 mL conical tube. The cultures were then incubated at 37°C with shaking for 3 h. To determine viable counts before antibiotic exposure, aliquots of each sorted group were serially diluted 10-fold in PBS and spot-plated (10 µL) onto LB agar plates as a control. Following ampicillin treatment, 900 µL of each culture was washed twice with PBS, resuspended in 30 µL of PBS, and spot-plated to quantify surviving cells. To confirm that sorting itself did not alter cell viability, two unsorted controls containing the same total number of cells (10^5^) were prepared. One control was serially diluted and plated prior to antibiotic treatment, while the second control was treated with ampicillin for 3 h under identical conditions. Survival of these unsorted controls was compared to that of the high-ribosome populations (groups C–D).

### Statistical analysis

The Student’s t test (two-sample assuming equal variances) was used to determine statistical significance using Microsoft Excel. *P* < 0.05 was considered significant. All experiments were conducted with at least duplicate replicates and were independently repeated at least twice.

## RESULTS & DISCUSSION

### GFP-based ribosome proxy reliably reflects intracellular ribosome levels

We previously demonstrated that *E. coli* MG1655-ASV, which carries an unstable GFP variant expressed from the chromosomal ribosomal promoter *rrnbP1* (Shah et al., 2006), serves as a reliable proxy for ribosome content. The *rrnbP1* promoter drives transcription of an operon encoding *rrsB* (16S rRNA), *gltT* (tRNA-glu), *rrlB* (23S rRNA) and *rrfB* (5S rRNA), thereby regulating the three major rRNA components of the ribosomes. Specifically, we showed GFP fluorescence in MG1655-ASV matches the abundance of active ribosomes during exponential growth (i.e., during balanced growth), rifampicin-induced dormancy (i.e., during persistence), and stationary-phase (i.e., during unbalanced growth) (Kim et al., 2018b). The ASV-tagged GFP variant has a half-life of less than 1 h, allowing real-time monitoring of ribosome synthesis dynamics. Moreover, previous studies have shown that rRNA abundance is a reliable surrogate for ribosome levels (Lu et al., 2009; Piques et al., 2009; Burger et al., 2010). Therefore, GFP intensity in MG1655-ASV was used as a quantitative proxy for ribosome content. A schematic overview of our approach, which examines how ribosome concentration influences persistence, is shown in **Fig. 1A**.

### Persistent cell formation is inversely proportional to ribosome level

To minimize the carryover of overnight-derived persister cells, cultures were inoculated at 0.5%. Any persisters present in the inoculum rapidly resuscitate (within minutes) upon transfer to fresh LB medium and regain high ribosome levels by the time the culture reaches an OD_600_ of 0.8 (Kim et al., 2018b). Using fluorescence-based gating, ∼10^5^ cells were sorted into four groups (A–D) spanning the full range of ribosome levels (**Fig. 2** and **Fig. S3**). As expected, a wide range of ribosomes per cell was found around a strong population average with some cells having very low ribosome levels (**Fig. 2**, Group A), which agrees with both our previous results while investigating ribosome levels during persister cell resuscitation (Kim et al., 2018b) and with single-cell ribosome results for *E. coli* growing exponentially on agarose pads (Chao et al., 2024); this heterogeneity in *E. coli* ribosome levels at the single cell level has also been described during different growth rates (Pavlou et al., 2025).

**Fig. 2.**
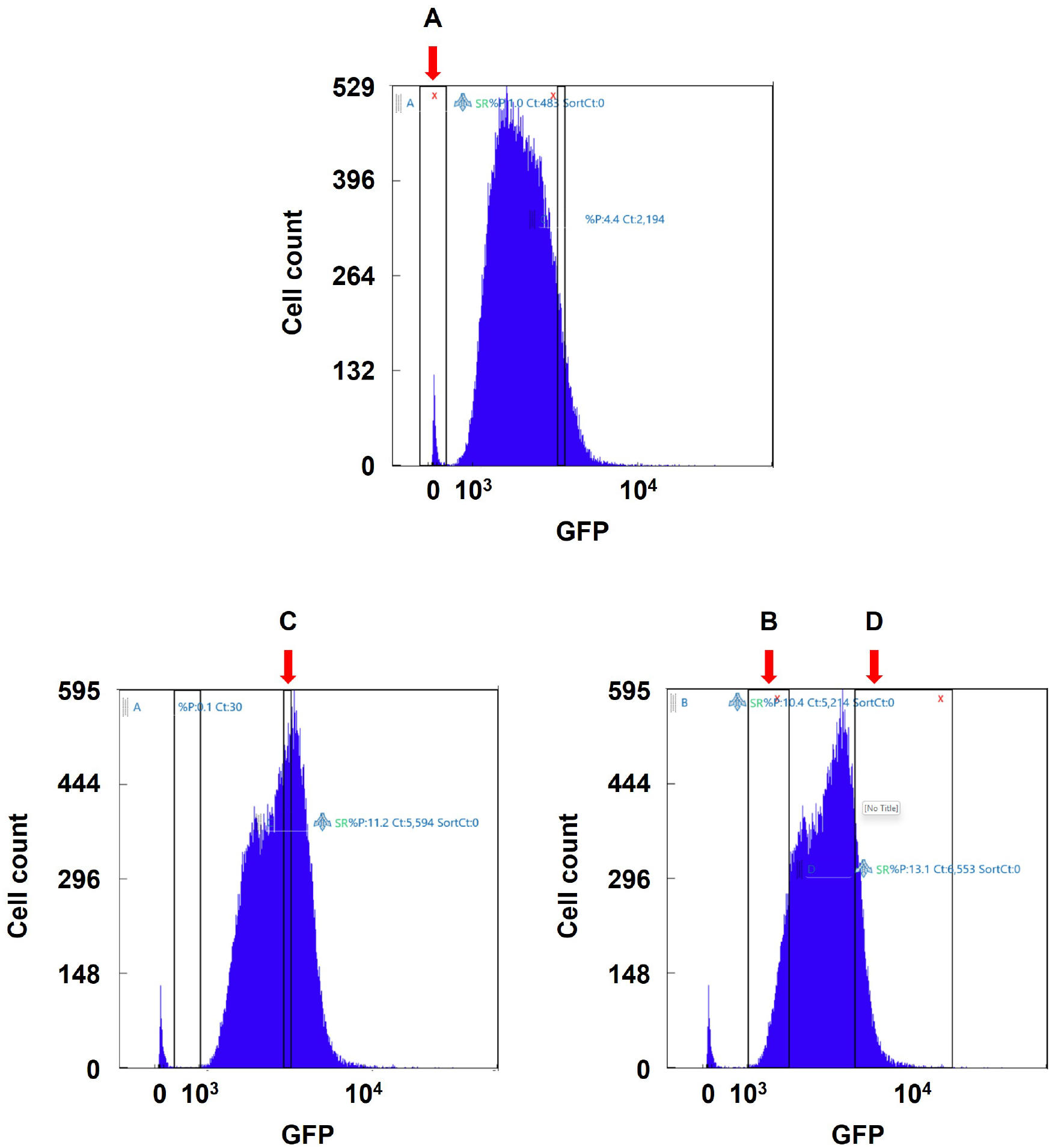
FACS sorting of cells based on GFP fluorescence intensity. Representative data for FACS sorting. Gates for Groups A, B, C, and D are indicated by arrows. Gates for the GFP-negative population (MG1655) are shown in **Fig. S1**.

Critically, cells with low ribosome content exhibited dramatically higher persistence: Group A, the lowest-ribosome subpopulation, showed an ∼80-fold increase in persister frequency relative to Group C, which reflects the average exponentially growing population (**Fig. 1B**; **Table 1**). As expected, Group B, which has substantially fewer ribosomes per cell (within the lowest 10% of the GFP-intensity distribution) than the population average (roughly equivalent to Group C), also exhibited a ∼3-fold increase in persistence compared to Group C.

**Table 1.** Comparison of persister levels among the four GFP-intensity–sorted groups (reflecting ribosome levels) following ampicillin treatment (3 h at 100 µg/mL,10× MIC). Data are presented as mean ± standard deviation, calculated from at least two independent experiments.

| Group | Condition | CFU/mL | Survival % | Fold change |
| --- | --- | --- | --- | --- |
| A | None | 300 ± 100 | 100 | 80 |
| | Amp | 4 ± 1 | 1.7 ± 0.5 | |
| B | None | 31,000 ± 3,000 | 100 | 3 |
| | Amp | 18 ± 2 | 0.063 ± 0.008 | |
| C | None | 30,000 ± 10,000 | 100 | 1 |
| | Amp | 8 ± 1 | 0.022 ± 0.004 | |
| D | None | 40,000 ± 20,000 | 100 | 1 |
| | Amp | 8 ± 1 | 0.029 ± 0.003 | |

Antibiotic survival (i.e., persistence) was ∼2% for Group A (the lowest-ribosome population) and ∼0.06% for Group B. In contrast, Group C, which contains the majority of the population, survived at only ∼0.02%. As expected, the survival of the Group C cells was similar to that of unfractionated exponentially growing cells (∼0.01% for GFP-positive cells) (**Table 2**), and this population-average survival also closely matched previously reported persister levels with ampicillin (0.01–0.2%) for *E. coli* BW25113 (Fernández-García et al., 2024). Groups C and D showed no detectable difference in either persister frequency or antibiotic survival (∼0.03%), despite Group D having even higher ribosome levels than Group C. Furthermore, GFP labeling did not affect persistence (0.01% for GFP-negative cells, **Table 2**). Together, these results indicate that neither GFP labeling nor the FACS sorting procedure affects cell survival and that higher ribosome levels than the population average do not alter persistence.

**Table 2.**
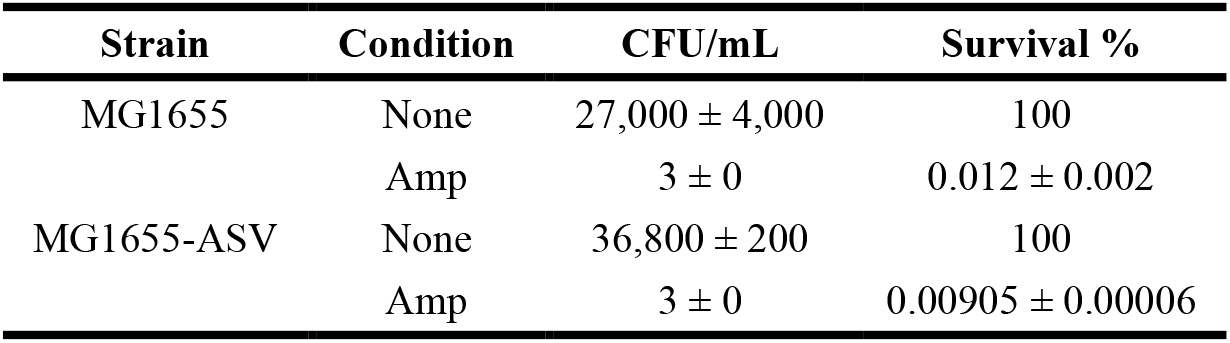
Comparison of persister levels between GFP-negative (MG1655) and GFP-positive (MG1655-ASV) strains prior to sorting, following ampicillin treatment (3 h at 100 µg/mL, ∼10× MIC). Data are presented as mean ± standard deviation, calculated from at least two independent experiments.

| Strain | Condition | CFU/mL | Survival % |
| --- | --- | --- | --- |
| MG1655 | None | 27,000 ± 4,000 | 100 |
| | Amp | 3 ± 0 | 0.012 ± 0.002 |
| MG1655-ASV | None | 36,800 ± 200 | 100 |
| | Amp | 3 ± 0 | 0.00905 ± 0.00006 |

### Ribosome abundance is a key determinant of persister formation

Our results show that ribosome content strongly governs the formation of persister cells. Here we find cells with low ribosome levels readily transition into the persister state (i.e., survive antibiotic treatment), whereas ribosome-rich cells rarely form persisters (**Fig. 1B**). Given that persister cells resuscitate rapidly based on ribosome content (Kim et al., 2018b), ribosome abundance functions as a central physiological determinant of both persister formation and resuscitation, and our findings provide strong support for the PRDP model. Moreover, since (i) persistence occurs in all bacterial and archaeal cells tested to date, (ii) cells primarily are undergoing stress (rather than exponential growth), and (iii) all lethal stress gives rise to the persister state, persistence may be the most important physiological state of cells, and our new results gives insights into how the persister state arises.

## SUSTAINABILITY STATEMENT

This review is most-related to sustainable goal 3, ‘Good health and wellbeing’, in that bacterial infection is likely to be the chief cause of death by 2050 (Udaondo and Matilla, 2020), and in order to treat these infections worldwide, persistence must be understood since persister cells, if not eradicated, may reconstitute these infections. Hence, our results are relevant to sustainability since they provide mechanistic insights into persister cell formation that should reduce mortality from bacterial infections.

## ACKNOWLEDGEMENTS

This work was supported by funds derived from NSF grant 2515448 (UEI: NPM2J7MSCF61). We are grateful for the assistance of Mitchell Koptchak and Rajeswaran Mani at the Pennsylvania State University Huck Institutes’ Flow Cytometry Core Facility (RRID:SCR_024460).

## DATA AVAILABILITY STATEMENT

All relevant data are contained within this manuscript.

## Supporting Information

**Fig. S1.**
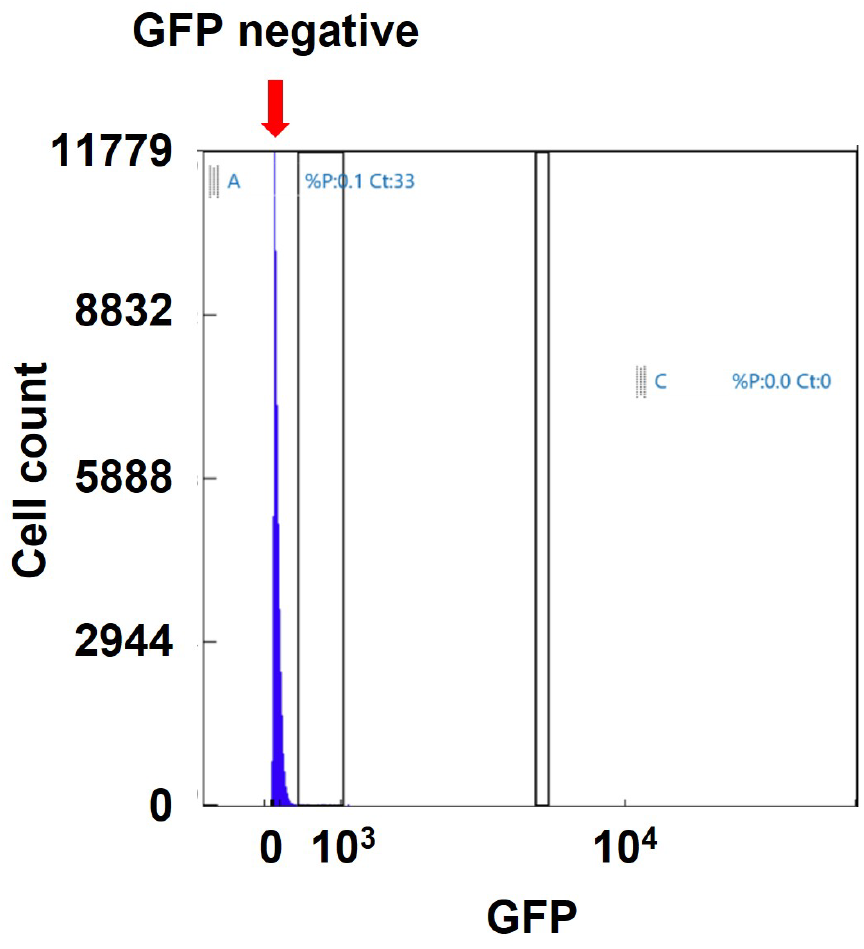
GFP fluorescence intensity of GFP-negative cells (representative data). MG1655, which lacks GFP, showed low-level fluorescence, as expected. The gate for the GFP-negative population, used to define the GFP-signal cut-off, is indicated by an arrow.

**Fig. S2.**
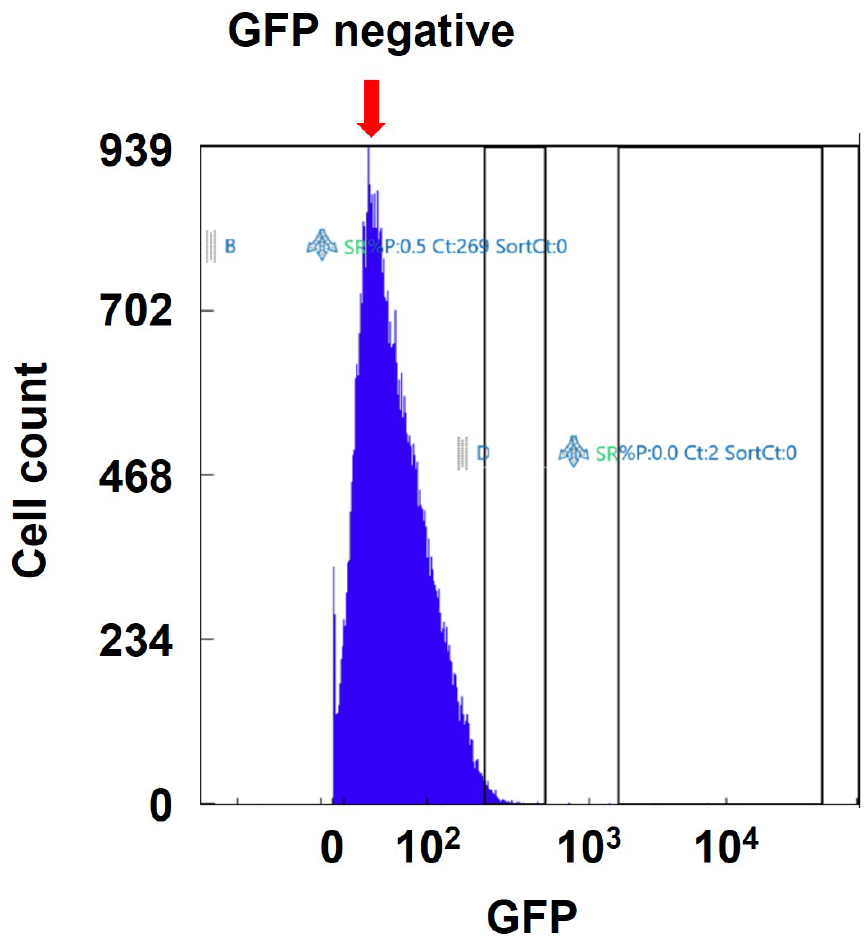
GFP fluorescence intensity of GFP-negative cells (data from an independent experiment). MG1655, which lacks GFP, showed low-level fluorescence, as expected. The gate for the GFP-negative population, used to define the GFP-signal cut-off, is indicated by an arrow.

**Fig. S3.**
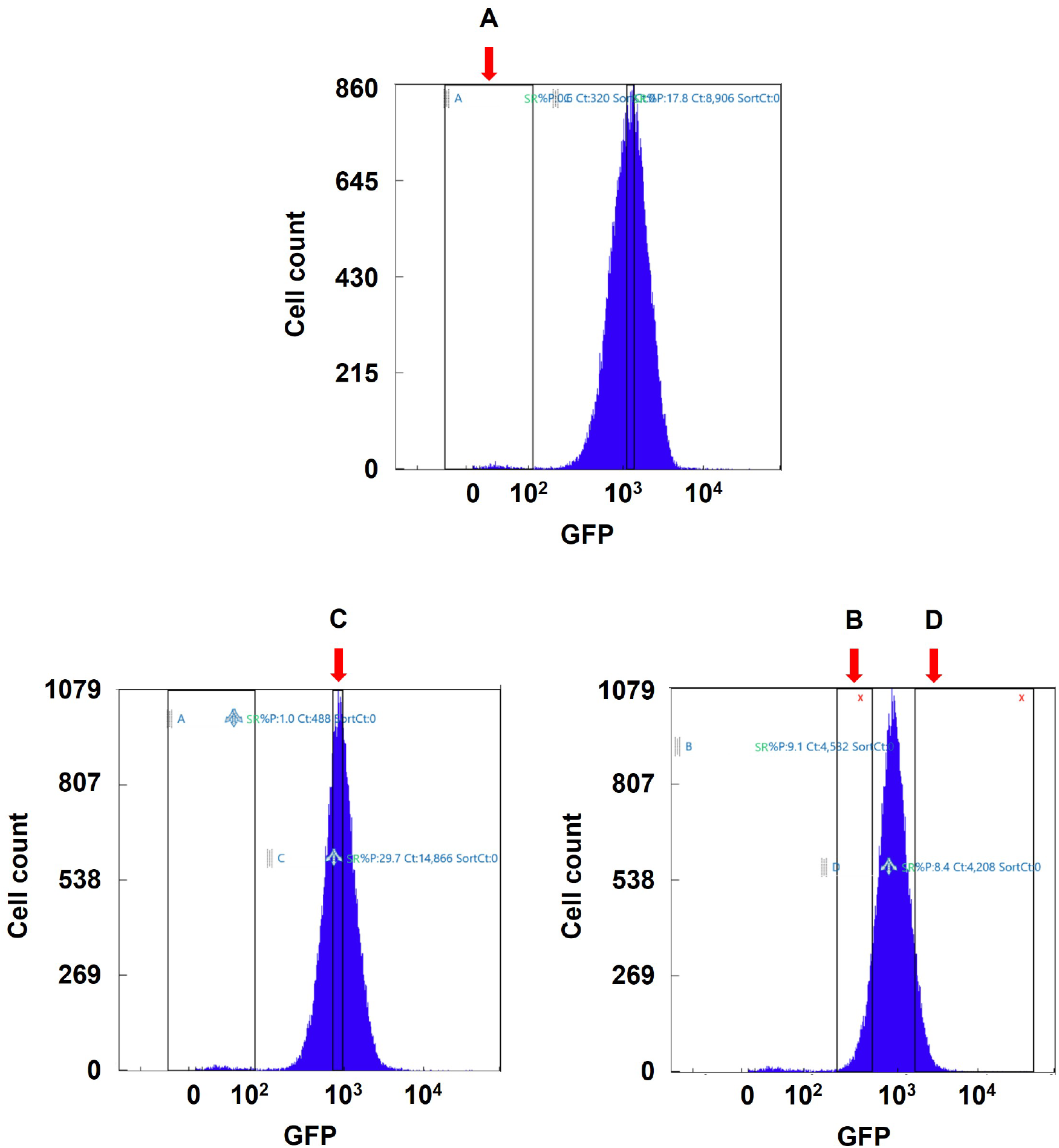
FACS sorting of cells based on GFP fluorescence intensity (data from an independent experiment). Data from an independent experiment showing FACS sorting. Gates for Groups A, B, C, and D are indicated by arrows. The GFP-negative population (MG1655) from this experiment is shown in **Fig. S2**.

